# *TP53* mutations drive therapy resistance via post-mitochondrial caspase blockade

**DOI:** 10.1101/2025.08.28.672283

**Authors:** Ahmed M. Mamdouh, Elyse A. Olesinski, Fang Qi Lim, Shaista Jasdanwala, Yang Mi, Nicole Si En Lin, Daniel Tan En Liang, Nishtha Chitkara, Leah Hogdal, R. Coleman Lindsley, Anthony Letai, Koji Itahana, Shruti Bhatt

## Abstract

Acute myeloid leukemia (AML) is a heterogeneous disease characterized by a broad spectrum of molecular alterations that influence clinical outcomes. *TP53* mutations define one of the most lethal subtypes of acute myeloid leukemia (AML), driving resistance to nearly all available treatment modalities, including venetoclax plus azacitidine (VenAza). Yet, the molecular basis of this resistance, beyond affecting transactivation of BCL-2 family genes, has remained elusive. Here, we demonstrate that VenAza treatment leads to reduced transcriptional upregulation of the p53 signaling pathway in *TP53* mutant/deficient AML compared to wild-type AML. Functionally, *TP53* mutant/deficient AML exhibits selective failure in apoptosis induction rather than impaired G1 arrest or senescence. Despite inhibition of pro-apoptotic BAX and selective enrichment for MCL-1 in *TP53* mutant isogenic AML cells, compensatory upregulation of BIM preserved functional mitochondrial outer membrane permeabilization (MOMP). *TP53* mutant primary AML tumors at baseline also had retained capacity for MOMP. Instead, *TP53* mutant AML exhibited disruption in caspase-3/7 activation to evade apoptosis after VenAza therapy – decoupling the mitochondrial and executioner phases of apoptosis. Importantly, this “post-MOMP brake” is not a bystander effect but itself a driver of VenAza and chemotherapy resistance in *TP53* mutant/deficient AML. This previously unrecognized mechanistic insight shifts the focus from mitochondrial priming to terminal caspase blockade in *TP53* mutant AML and opens the door for urgently needed therapeutic strategies that reignite apoptosis at its execution point.

## INTRODUCTION

*TP53* mutations or deletions are well-established drivers of therapy resistance across many cancers, leading to poor patient outcomes following chemotherapy and targeted therapy.^1–3^ In acute myeloid leukemia (AML), an aggressive hematological malignancy, *TP53* alterations confer particularly dismal prognoses. These mutations occur in ∼10% of *de novo* AML cases, 20-35% of therapy-related myeloid neoplasia, and up to 70-80% of complex karyotype AML (CK-AML).^4–6^ The advent of venetoclax (BCL-2 inhibitor) combined with azacitidine (hypomethylating agent) (VenAza) has transformed outcomes for many AML patients^7^, including those with poor-risk cytogenetics and wild-type *TP53,* who traditionally responded poorly to 7+3 intensive chemotherapy.^8–11^ However, VenAza offers very little benefit to patients with *TP53* mutations, particularly when co-occurring with complex karyotypes. In this subgroup, median overall survival plummets from 23.4 months in wild-type cases to just 5.2 months in *TP53* mutant patients.^7^ This stark disparity persists regardless of cytogenetic risk status, underscoring *TP53* mutations as an independent predictor of treatment failure. Patients harboring *TP53* mutations also experience significantly shorter relapse-free survival (4.3 months vs 18.9 months) and overall survival (5.2 months vs 19.4 months), emphasizing the urgent need for new therapeutic strategies.^11–13^

The tumor suppressor p53 functions as the “guardian of the genome” by transactivating genes in response to stress, including those regulating genomic integrity, cell cycle control, and apoptosis induction.^11,14^ Canonically, p53 induces apoptosis primarily by triggering mitochondrial outer membrane permeabilization (MOMP) via transcriptional activation of pro-apoptotic BCL-2 family members (e.g. BAX, PUMA, and NOXA).^15,16^ Supporting this, studies have shown that *TP53* null cells cannot activate BAX following venetoclax treatment to induce MOMP.^17^ Additionally, p53 can be sequestered in the cytoplasm by anti-apoptotic BCL-2 proteins (e.g. MCL-1, BCL-2 and BCL-xL) and released by PUMA to promote MOMP.^18–22^ While MOMP is considered the ‘point-of-no-return’ in apoptotic signaling, emerging studies suggests that even after MOMP, wild-type p53-expressing cells may recover^23^, indicating the existence of additional regulatory steps downstream of mitochondrial permeabilization.

Despite p53’s central role in apoptosis, the precise mechanisms by which *TP53* mutations drive therapy resistance remain incompletely defined. Here, we identify among different functions of p53 that disruption in apoptosis is the leading contributor to VenAza resistance. One may expect evasion of apoptosis due to failure of MOMP induction in *TP53* mutant/null cells, but we show that MOMP remains proficient. Instead, *TP53* mutant AML escapes apoptosis via failure to activate post-mitochondrial executioner caspases. These novel findings highlight that *TP53* mutant AML effectively uncouples mitochondrial permeabilization from post-mitochondrial signaling to evade apoptosis.

## MATERIALS AND METHODS

### Alkaline comet assay

We pre-coated glass slides with 1% (w/v) agarose in type I grade water and dried them overnight at room temperature. For each treatment group, 1.0 x 10^5^ cells were resuspended in 100 µL fresh media and 200 μL of 1% (w/v) low-melting agarose in PBS. Samples were added to pre-coated slides and cooled at 4°C to solidify. Slides were treated with alkaline lysis buffer by immersion in an alkaline electrophoresis solution at 4°C for 40 minutes before electrophoresis was conducted at room temperature. Slides were neutralized, fixed, stained with propidium iodide, and rinsed with PBS. Images were collected on a Nikon SMZ25 fluorescence microscope. Quantification done using CometScore 2.0.

### Beta-galactosidase senescence assay

1.0×10^6^ cells were treated before analysis by flow cytometry according to the manufacturer’s protocol (Invitrogen). Cells were fixed in 4% paraformaldehyde for 10 minutes at room temperature. Subsequently, cells were stained with CellEvent Senescence Green Probe, diluted at a ratio of 1:2000 in CellEvent Senescence Buffer, and incubated at 37°C in the dark for 1 hour. Data was acquired on Attune NxT Flow Cytometer (Invitrogen) and analyzed on FlowJo. For positive control, U2OS cells were treated with nocodazole (100 ng/mL) for 96 hours.

### BH3 profiling and Dynamic BH3 profiling

To determine priming for apoptosis, 2.5×10^4^ cells were resuspended into 15 µL of MEB buffer and seeded into a 384-well plate. BIM peptide (determinant for overall priming) in different concentrations was mixed with 0.04% digitonin and added into cells. DMSO and alamethicin were used as negative and positive controls, depicting 100% and 0% cytochrome c retention, respectively. Mitochondria of permeabilized cells were exposed to peptides for 1 hour at room temperature with shaking and fixed in the dark with 4% paraformaldehyde for 10 minutes with shaking. Cells were subsequently neutralized with N2 buffer for 15 minutes with shaking and stained with Hoescht and anti-cytochrome c-Alexa647 (BioLegend, # 612310). At 24 hours post-incubation, quantification of cytochrome c release induced by each peptide was analyzed using flow cytometry (FACSCelesta). Mitochondrial priming was measured as FACS-determined cytochrome c loss = 100 − (% of cells within cytochrome c retention gate).^24^ For dynamic BH3 profiling, cells were treated with the desired agent for 4 hours before harvesting, after which samples were processed in the manner mentioned above. Drug-induced change in priming, or “delta priming,” was calculated as = cytochrome c loss [drug] – cytochrome c loss [DMSO].

### Caspase luminescence assay

1.0×10^4^ MOLM-13 cells were seeded in 100 µL RPMI per well in 96-well plates and treated with agents of interest. After 24 hours of incubation at 37°C, 30 μL was aliquoted from each well and added to a 384-well plate. Next, 30 μL of Caspase Glo (Promega) was added and incubated for 1.5 hours at room temperature. The luminescence intensity was measured by a Hidex sense multi-plate reader.

### Cell culture

AML cell lines were obtained as follows: MOLM-13 from DSMZ, MV-4-11 from ATCC. MOLM-13-*TP53* isogenic AML cell lines *TP53^+/+^* (wild-type), *TP53^-/-^* (knock-out), and 6 mutants, *TP53^-/R248Q^*, *TP53^-/Y220C^*, *TP53^-/R82W^*, *TP53^-/R173H^*, *TP53^-/R275H^*, and *TP53^-/M237I^* were kindly provided by Steffen Boettcher.^25^ Lenti-X 293T, and U2OS cell lines were kindly provided by Nicholas Gascoigne, and Karen Crasta, respectively. Isogenic MOLM-13 cells were cultured in RPMI-1640 (Gibco, 11875093). Lenti-X 293T, and U2OS cells were cultured in DMEM (Biowest, L0104). Media was supplemented with 10% fetal bovine serum (Gibco, 10437028) and 1% penicillin-streptomycin (Gibco, 15140122). All cell lines were cultured in a humidified incubator at 37°C with 5% CO_2_.

### Cell cycle and cell death studies

To determine cell cycle changes within a cell line, 5.0×10^5^ cells were permeabilized and fixed using 70% ice cold ethanol and maintained at 4°C. Next, cells were washed with PBS, stained with 400 µL of DAPI with 0.1% Triton-X, and incubated for 20 minutes. Fixed cells were analyzed using flow cytometry (LSRFortessa X-20) at an ultraviolet wavelength of 496 nM. To measure cell death, triplicates of 1.0×10^4^ cells were seeded in the 96-well plate with either DMSO (control) or drug of interest and incubated at 37°C for treatment duration. Cells were stained with annexin V and propidium iodide solutions 10 minutes prior to acquisition on Attune NxT Flow Cytometer. Data was analyzed on FlowJo.

### Cytotoxicity assay

To determine viability of cells, 1.0×10^4^ cells were treated with corresponding agents at various concentrations in 96-well plates and incubated for 72 hours at 37°C. CellTiter-Glo (Promega) was added in a volume ratio of 4:1 (cell culture media:work solution), vigorously shaken for 2 minutes, and incubated for 10 minutes at room temperature. Plates were protected from light before the acquisition of luminescence signaling using a Hidex sense multi-plate reader. Absolute viability was converted to percentage viability vs. DMSO control treatment. Subsequently, IC50 values and a nonlinear fit of log (inhibitor) vs. normalized response was obtained using GraphPad Prism.

### Generation of knockout cells

sgRNA sequences were cloned into lentiCRISPRv2 (Addgene, #52961). Lenti-X 293T cells (1.8×10^6^ cells) were transfected with envelope plasmid pVSV-G (415 ng), packaging plasmid psPAX2 (830 ng), and the corresponding transfer plasmids (1245 ng) using Lipofectamine™ 3000 transfection reagent (ThermoFisher, L3000015) and Opti-MEM (Gibco, 31985070).

Cell culture media was replaced with DMEM after 24 hours. Supernatants containing lentiviruses were collected at 48 hours and 72 hours post transfection and filtered through a 0.45 μM nitrocellulose filter (Sartorius). AML cells were centrifuged at a speed of 2500 rpm for 90 minutes at 32°C with 200 μL of virus-containing media supplemented with 10 μg/mL polybrene (MedChemExpress). Puromycin (2 μg/mL) or hygromycin (400 µg/ml) was added 48 hours post-infection. sgRNA sequences are listed in Supplemental Table 1.

### Statistical significance

Calculation of statistical significance was conducted using two-tailed t-tests unless otherwise specified. For all figures, *p<0.05, **p<0.01, ***p<0.001, ****p<0.0001.

### TMRE staining

Cells were seeded at a density of 1.0×10^6^ cells per mL of media. The cells were stained with TMRE (10 nM) (Invitrogen) and incubated at 37°C with 5% CO_2_ for 30 minutes. Cells were pelleted before resuspending in fresh media and treated. Cells were further incubated for 24 hours before staining with DAPI and analysis by flow cytometry.

### Western blot and coimmunoprecipitation

Cells were lysed in NP-40 lysis buffer (150 mM NaCl, 1% NP-40, 50 mM Tris-Cl pH 8.0) for coimmunoprecipitation or cell lysis buffer (Cell Signaling Technology #9803) for western blot supplemented with protease inhibitor and phosphatase inhibitor (when needed) for 45 minutes at 4°C according to the manufacturer’s protocol. Mitochondria were isolated using the Mitochondria Isolation Kit (ThermoFisher). Protein concentration was determined using Pierce BCA Protein Assay Kit (ThermoFisher). For western blot: Proteins were boiled in Laemmli Sample Buffer (Bio-Rad) containing 10% β-mercaptoethanol (Sigma-Aldrich), separated by SDS-PAGE, and transferred onto nitrocellulose membranes. For coimmunoprecipitation: 300 μg of pre-cleared lysates from each sample was incubated with Dynabeads Protein G beads (ThermoFisher) overnight prior to incubation with BIM antibody (Cell Signaling Technology, #2933) at 4°C for 4 hours. Beads were washed with NP-40 lysis buffer and washing buffer before eluting using elution buffer from Dynabeads™ Protein G Immunoprecipitation Kit (ThermoFisher). For both: Membranes were blocked for 1 hour with 5% milk (Bio-Rad) dissolved in TBST (w/v) and incubated overnight in primary antibodies diluted in 5% milk. On the next day, membranes washed with TBST (w/v) and incubated with secondary antibody for 1 hour, then washed again with TBST (w/v) before developing the plots using SuperSignal West Femto Maximum Sensitivity Substrate (ThermoFischer). List of antibodies used are listed in Supplemental Table 2. For comparison between blots in Figure 6C and S6A, images were acquired simultaneously.

## RESULTS

### *TP53* mutation/deletion causes resistance to VenAza

Although the clinical challenge of therapy resistance in *TP53* mutant AML has been recognized, the underlying mechanisms remain incompletely understood. Hence to study the mechanisms of resistance, we engineered *TP53* deficient AML cells using CRISPR/Cas9 genome editing in MOLM-13, and MV-4-11 with two different sgRNAs (Figures 1A and S1A). Successful knock-out was validated by evaluating the expression of *TP53* and *CDKN1A* (Figure S1B). *TP53* deletion reduced sensitivity to venetoclax plus azacytidine (VenAza) compared to wild-type *TP53* (Figure 1B). In addition to deletions, *TP53* dysfunction is also caused by missense mutations that cause variable p53 expression (Figure S1C). Thus, we exposed a panel of isogenic MOLM-13 cells carrying six different missense mutations *TP53^-/mut^* (*R175H, R273H, R248Q, R282W, Y220C, M237I*)) in the DNA binding domain (DBD), knockout (*TP53^-/-^*), or matched wild-type (*TP53^+/+^*) to venetoclax-only and azacitidine-only. All *TP53^-/mut^* and *TP53^-/-^* had less sensitivity compared to *TP53^+/+^* when treated with either venetoclax or azacitidine as a single agent, as shown before^26^ (Figure 1C). To ask if there is differential sensitivity among the six distinct *TP53* missense mutations, we combined low concentration azacitidine (50 nM) with venetoclax. Indeed, *TP53^-/mut^* cells had varied sensitivities to VenAza, with *TP53^-/R248Q^*cells the least sensitive (IC50 for VenAza combination = 72.5 nM) (Figure 1D). The *R248Q* mutation is also second most prevalent *TP53* mutation among other missense mutations in myeloid malignancies.^25^ For this reason, we decided to focus on the R248Q mutation. *TP53^-/R248Q^* cells also had higher colony forming units (CFUs) under VenAza compared to *TP53^+/+^* (76.9% vs 56.4%; Figure 1E). All 6 *TP53^-/mut^* and *TP53^-/-^* cell lines had a competitive advantage over *TP53^+/+^* cells treated with VenAza (Figures 1F and S1D). In conclusion, *TP53* mutation or deletion causes resistance to Ven alone, Aza alone, or VenAza combination.

**Figure 1.**
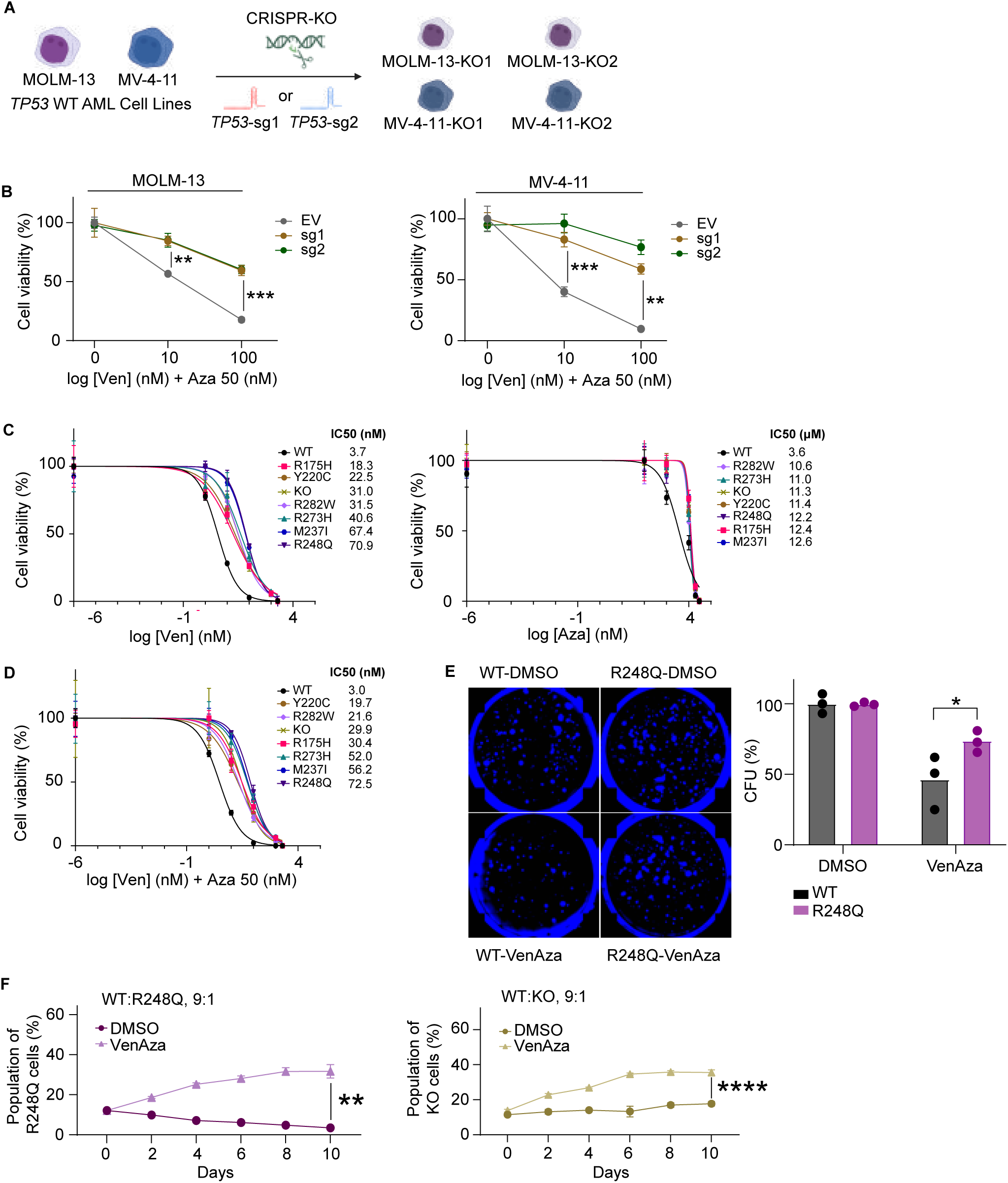
*TP53* mutant AML is resistant to VenAza. **(A)** Schematic of CRISPR-KO generation of *TP53* knockout MOLM-13. **(B)** Cell viability of *TP53* active (EV) and knockout (sg1, sg2) AML cells (MOLM-13, MV-4-11) treated with venetoclax and azacitidine. **(C)** Cell viability of isogenic MOLM-13 cells after 72 hours of treatment with increasing concentrations of venetoclax or azacitidine as a single agent. **(D)** Cell viability of isogenic MOLM-13 cells after 72 hours of treatment with increasing concentrations of venetoclax and azacitidine. **(E)** Representative images of colony formation assays after treatment with DMSO or VenAza for 72 hours in RPMI followed by 9 days in Methocult. Colony forming units were normalized to respective DMSO treated cells. **(F)** Clonal competition assay evaluating the competition between the cells in co-culture after treatment with DMSO or VenAza for 10 days. For all figures unless otherwise specified, treatment with VenAza: venetoclax 3 nM, azacitidine 50 nM. * p<0.05, ** p<0.01, *** p<0.001, **** p<0.0001.

### TP53 mutations impair VenAza-induced p53 signaling and associated transcriptional programs

Since p53 coordinates key stress response pathways, we investigated whether VenAza response is affected in *TP53* genotypes via transcriptional regulation of its canonical functions – DNA damage response, cell cycle arrest, and apoptosis – which has previously been evaluated in *TP53^-/-^* lymphoma^27^ but not AML. To this end, we performed RNA sequencing on isogenic MOLM-13 cells harboring *TP53^+/+^*, *TP53^-/R248Q^*, or *TP53^-/-^* alleles following treatment with VenAza or DMSO as a control (Figure 2A). Principal component analysis (PCA) revealed clear segregation of transcriptional signatures by *TP53* genotype (Figures 2B and S2A). Further, VenAza exposure also modulated distinct transcriptional programs within each genotype.

**Figure 2.**
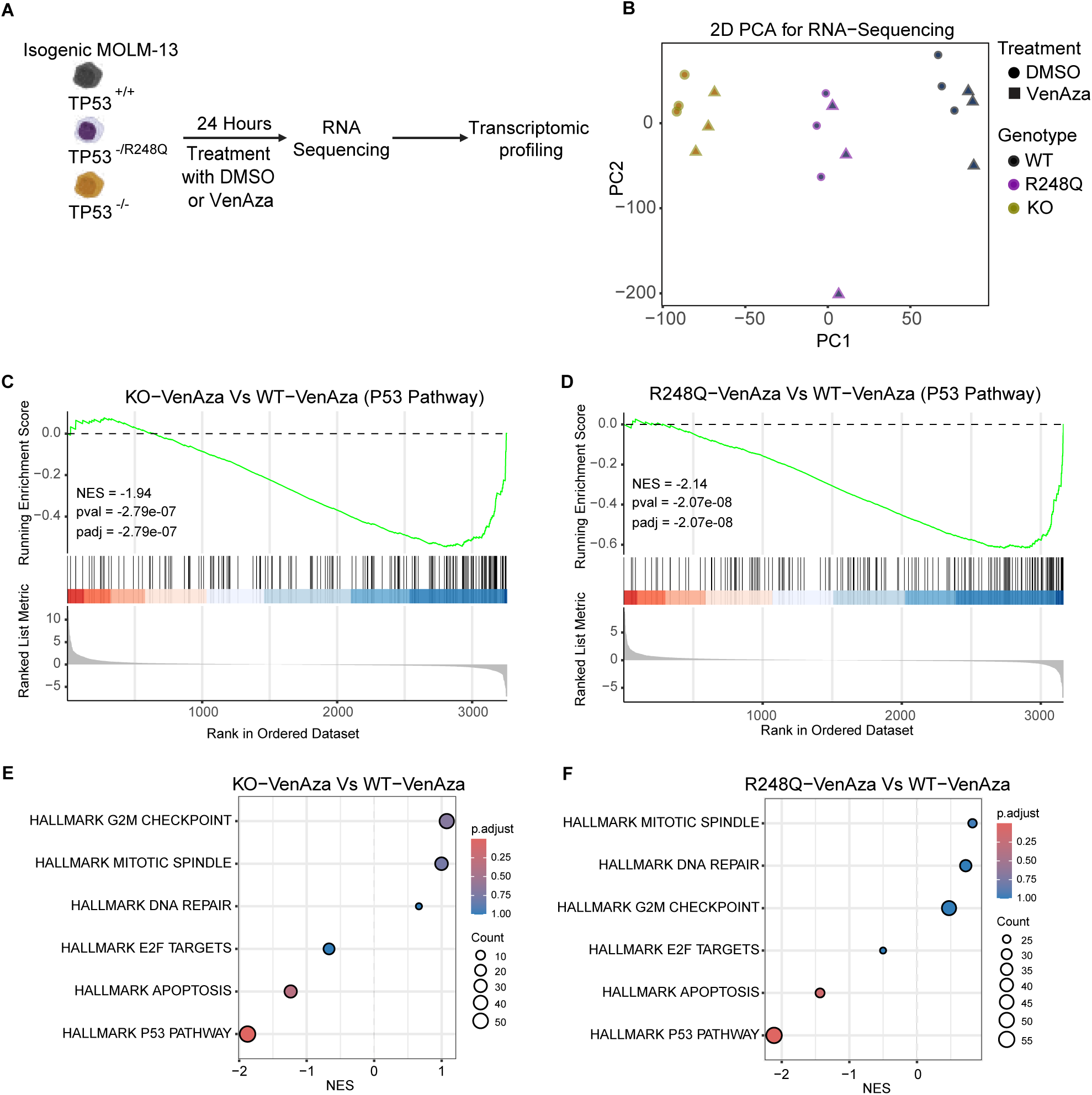
Transcriptomic profiles of *TP53* mutant AML. **(A)** Schematic overview of the experimental design of RNA sequencing. Isogenic MOLM-13 cells were treated with DMSO or VenAza for 24 hours, followed by RNA-seq for transcriptomic profiling. **(B)** Principal component analysis (PCA) of RNA-seq data from all samples. PC1 and PC2 distinguish samples based on genotype and treatment. Genotypes are color-coded, while treatment conditions are indicated by shape. **(C-F)** Gene set enrichment analysis (GSEA) using MsigDB for 6 hallmarks related to the main p53 canonical functions: **(C)** p53 pathway for *TP53^-/-^* treated with VenAza vs *TP53*^+/+^ treated with VenAza; **(D)** p53 pathway for *TP53^-/R248Q^* treated with VenAza vs *TP53*^+/+^ treated with VenAza; **(E)** other pathways for *TP53^-/-^* treated with VenAza vs *TP53*^+/+^ treated with VenAza; **(F)** other pathways for *TP53^-/R248Q^* treated with VenAza vs *TP53*^+/+^ treated with VenAza. For all figures unless otherwise specified, treatment with VenAza: venetoclax 3 nM, azacitidine 50 nM.

Hence, we next conducted gene set enrichment analysis (GSEA) using the MsigDB Hallmark collection to evaluate gene sets for the p53 pathway, DNA repair, apoptosis, and cell cycle (E2F targets for G1-S phase, G2M checkpoint, and mitotic spindle). As anticipated, p53 signaling was down-regulated in *TP53^-/R248Q^* and *TP53^-/-^* cells at baseline compared to *TP53^+/+^* cells (p < 0.0001; Figures S2B-C). We then asked whether VenAza exposure influenced p53 signaling in a genotype-specific manner. Compared to DMSO, VenAza led to significant enrichment in p53 pathways for *TP53^+/+^* (p < 0.0001) and *TP53^-/R248Q^*(p < 0.05) cells but not *TP53^-/-^* (p > 0.05) cells (Figures S2D-F). Remarkably, the induction of p53 signaling was significantly suppressed in *TP53^-/R248Q^* and *TP53^-/-^* cells compared to *TP53^+/+^* cells following VenAza treatment (p < 0.0001; Figures 2C-D). Among other p53-regulated canonical pathways, the apoptosis pathway was modestly downregulated, while DNA repair, G2M, and mitotic spindle pathways were modestly upregulated (p > 0.05; Figures 2E-F).

Together, this indicates that intact p53 signaling in AML in response to VenAza is upregulated in *TP53^+/+^* cells compared to *TP53^-/R248Q^* and *TP53^-/-^* cells. Nevertheless, programs involved in cell cycle progression and DNA repair are less affected transcriptionally in response to VenAza.

### *TP53* mutations selectively impair VenAza-induced apoptosis without affecting cell cycle progression or senescence

After transcriptomic profiling, we next investigated whether VenAza treatment functionally affects *TP53*’s main canonical roles in cell cycle regulation, apoptosis, senescence, and DNA damage repair. To test this, we treated isogenic MOLM-13 cells carrying six different *TP53* DNA binding domain missense mutations (*TP53^-/mut^: R175H, R273H, R248Q, R282W, Y220C, M237I*), *TP53* knockout (*TP53^-/-^*), or matched *TP53* wild-type (*TP53^+/+^*) with VenAza. At baseline, *TP53^-/R248Q^*and *TP53^-/-^* cells showed no major differences in cell cycle profiles or senescence compared to *TP53^+/+^*cells (Figures 3A and S3A-B). However, following VenAza, *TP53^-/R248Q^* and *TP53^-/-^* cells failed to accumulate in the sub-G1 phase, indicating defective apoptosis despite intact mitotic entry that supported transcriptomic changes. Apoptosis was severely blunted across all *TP53^-/mut^* and *TP53^-/-^* MOLM-13 cells in response to both VenAza and standard chemotherapy (cytarabine or etoposide) and isogenic *TP53^-/-^* MV-4-11 cells treated with VenAza (Figures 3B-C).

**Figure 3.**
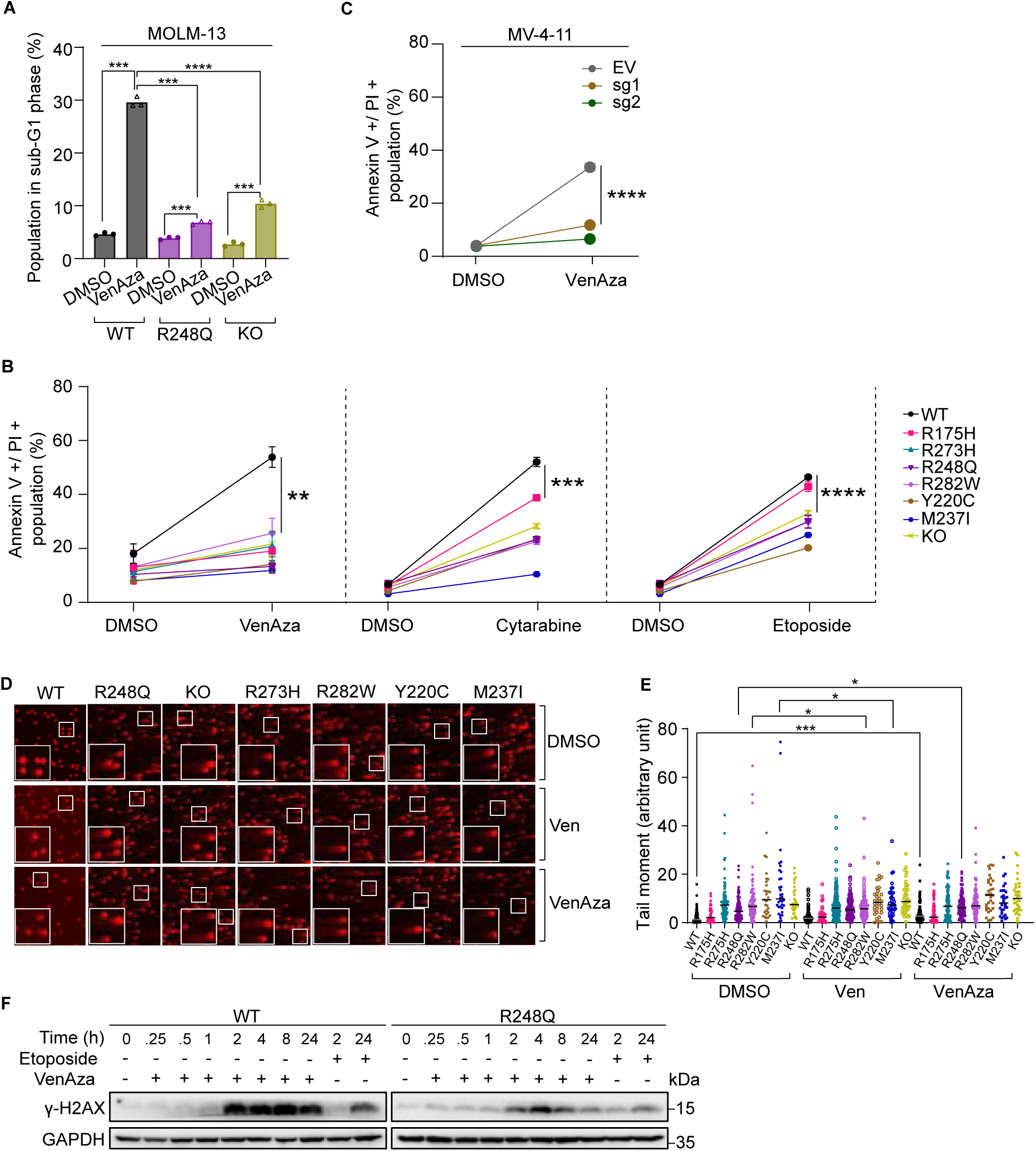
*TP53* mutant AML resists VenAza treatment by evasion of apoptosis. **(A)** Population of cells in sub-G1 phase MOLM-13 *TP53*^+/+^, *TP53*^-/R248Q^, and *TP53^-/-^* cells treated with DMSO or VenAza for 24 hours. **(B)** Annexin V and PI assay demonstrate greater viable cell populations in *TP53^-/mut^* and *TP53^-/-^* isogenic MOLM-13 cells treated with VenAza, cytarabine (200 nM), or etoposide (200 nM) for 72 hours. **(C)** Annexin V and PI assay for MV-4-11 after treatment with venetoclax 10 nM plus azacitidine 50 nM for 72 hours. **(D)** Alkaline comet assay of isogenic *TP53^+/+^, TP53^-/mut^*, *TP53^-/-^* cells MOLM-13 cells treated with DMSO, venetoclax (1 nM), or VenAza (venetoclax 1 nM, azacitidine 50 nM) for 7 days. **(E)** Quantification of comet tails normalized to *TP53*^+/+^ DMSO. **(F)** Time course western blot of γ-H2AX after treatment with VenAza or etoposide (0.5 uM). For all figures unless otherwise specified, treatment with VenAza: venetoclax 3 nM, azacitidine 50 nM. * p<0.05, ** p<0.01, *** p<0.001, **** p<0.0001.

To assess whether this apoptotic defect was linked to DNA damage, we performed alkaline comet assays. In *TP53^+/+^* cells, VenAza increased DNA damage relative to DMSO (Figures 3D-E). In contrast, *TP53^-/mut^* cells displayed elevated baseline DNA damage that did not further increase with treatment. We also assessed γ-H2AX levels as a marker of double-strand DNA breaks. *TP53^+/+^* cells showed rapid induction of γ-H2AX following treatment, with levels surpassing those observed in *TP53^-/R248Q^*cells at both early and late time points (Figure 3F). These results suggest that *TP53* mutations drive resistance to cytotoxic and targeted therapies in AML by disrupting apoptosis and resisting DNA damage effects, rather than through disruption of senescence or cell cycle signaling.

### *TP53* mutations paradoxically upregulate BIM while enhancing its sequestration by MCL-1

To investigate mechanisms of apoptotic resistance in *TP53* mutant AML, we next examined expression of BCL-2 family proteins – key regulators of the intrinsic mitochondrial pathway and therapeutic targets of venetoclax. As expected, *TP53* target genes *CDKN1A* and *MDM2* transcripts were downregulated in *TP53^-/R248Q^* and *TP53^-/-^* cells at baseline (Figure S4A). Since *TP53* regulates apoptotic genes, we hypothesized that VenAza would alter expression of BCL-2 family genes and proteins in a *TP53*-dependent manner. At the mRNA level, pro-apoptotic genes *BAX*, *BBC3* (encodes PUMA), and *PMAIP1* (encodes NOXA) were downregulated in *TP53^-/R248Q^* cells compared to compared to *TP53^+/+^* cells at the baseline as well as in the presence of VenAza (Figure S4A). At the protein level, pro-apoptotic protein BAX was decreased, and the anti-apoptotic protein MCL-1 was increased (Figures 4A and S4B). Conversely, pro-apoptotic proteins BIM-EL, BAK, and BID were upregulated. Since BIM can directly interact with and activate BAX and BAK (required for pore formation and MOMP), we further investigated BIM expression levels across multiple tumors. MV-4-11 cells showed upregulation in BIM-EL after *TP53* deletion (Figure 4B and S4C). Similarly, primary tumors from the TCGA Pan-Cancer Atlas showed upregulation in BIM for 5/27 tumors at the mRNA level and 10/26 at the protein level (Figures 4C and S4D). This emphasizes that despite loss of key pro-apoptotic regulators in *TP53* mutations, upregulation of BIM and BID may compensate for this downregulation.

**Figure 4.**
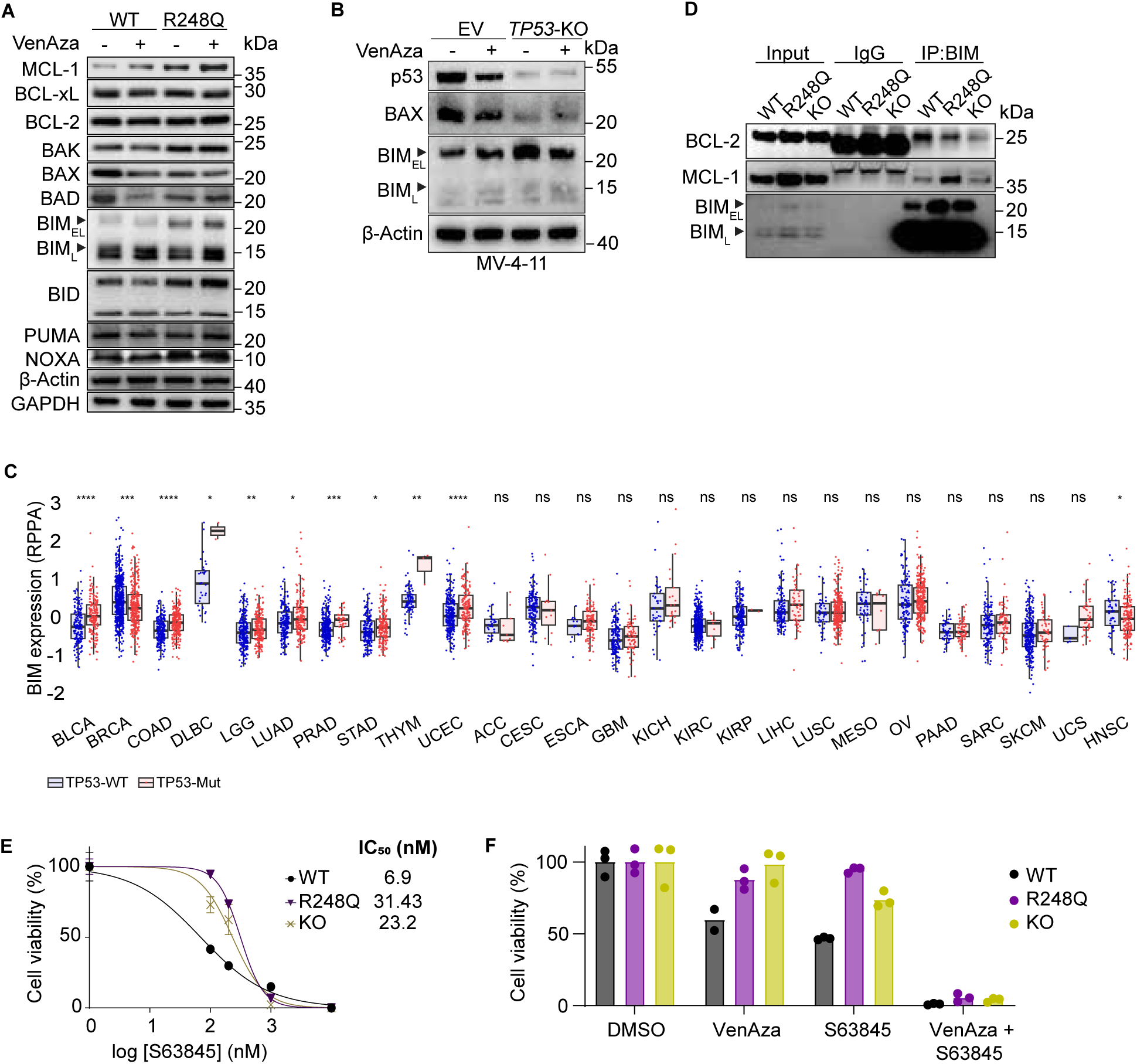
*TP53* mutations alter BCL-2 family proteins and respond to simultaneous BCL-2/MCL-1 targeting. **(A)** Immunoblotting of BH3 family proteins and other apoptotic factors in *TP53^+/+^* and *TP53^-/R248Q^* cells treated with DMSO or VenAza for 24 hours. **(B)** Western blot of *EV* and *TP53^-/-^*MV-4-11 cells treated with DMSO or VenAza (venetoclax 10 nM, azacitidine 50 nM) for 24 hours. **(C)** BIM protein expression levels across multiple cancers from the TCGA database, stratified according to *TP53* status. Analysis via Wilcoxon test. **(D)** Coimmunoprecipitation assay identifies interactions between BIM:MCL-1 or BIM:BCL-2. **(E)** Cell viability of MOLM-13 isogenic cells after exposure to **(E)** S63845 for 72 hours or **(F)** DMSO, VenAza, S63845 (10 nM), or VenAza plus S63845 (10 nM) for 72 hours. For all figures unless otherwise specified, treatment with VenAza: venetoclax 3 nM, azacitidine 50 nM. * p<0.05, ** p<0.01, *** p<0.001, **** p<0.0001.

Sequestration of BIM by anti-apoptotic BCL-2 family members can inhibit apoptosis. Prior studies, including ours, have shown that in venetoclax-resistant AML, MCL-1 sequesters BIM away from BCL-2.^28–30^ Coimmunoprecipitation assays revealed reduced BCL-2–BIM interactions and increased MCL-1–BIM interactions in these cells (Figure 4D and S4E). Although *TP53^-/R248Q^*and *TP53^-/-^* cells were more resistant to MCL-1 inhibitor S63845; combining the MCL-1 inhibitor with VenAza restored sensitivity in *TP53* mutant cells, consistent with prior reports^31^ (Figures 4E-F). However, while targeting MCL-1 is a rational approach to overcome resistance, clinical development has been hampered by dose-limiting cardiotoxicity.^32^ Together, *TP53* mutations reprogram both expression and functional interactions of BCL-2 family proteins.

### MOMP occurs in *TP53* mutant AML despite alternations in BCL-2 family proteins

Apoptosis critically depends on MOMP, a key step in the intrinsic apoptotic pathway often considered the ‘point of no return.’ Thus, we hypothesized that *TP53* mutant cells evade apoptosis by resisting MOMP induction. We performed BH3 profiling assays on unperturbed, permeabilized *TP53^-/mut^* cells using BH3 peptides derived from pro-apoptotic proteins. Across multiple contexts – *TP53^+/+^*, *TP53^-/mut^*, and *TP53^-/-^* isogenic MOLM-13 cells and *TP53^-/-^* MOLM-13 and MV4-11 edited with distinct *TP53* sgRNAs – we consistently observed retained capacity for MOMP induction in response to the BIM BH3 peptide (Figures 5A and S5A). Notably, these findings suggest that *TP53* mutant and deficient cells maintain intact apoptotic priming. Further, BH3 profiling revealed similar dependencies on individual anti-apoptotic proteins – BCL-2, (BAD BH3 peptide), MCL-1 (MS-1 BH3 peptide), or BCL-xL (HRK BH3 peptide) – in *TP53^-/mut^*and *TP53^+/+^* cells (Figure S5B). Primary AML samples showed comparable BIM-induced priming across *TP53* statuses and preserved mitochondrial sensitivity to navitoclax and venetoclax. Increased dependence on BCL-xL and MCL-1 was also observed, as indicated by response to HRK and NOXA peptides, respectively (Figures 5B-C). Lastly, preserved MOMP in *TP53* mutant cells was not attributable to differences in baseline cytoplasmic cytochrome c levels (Figure S5C).

**Figure 5.**
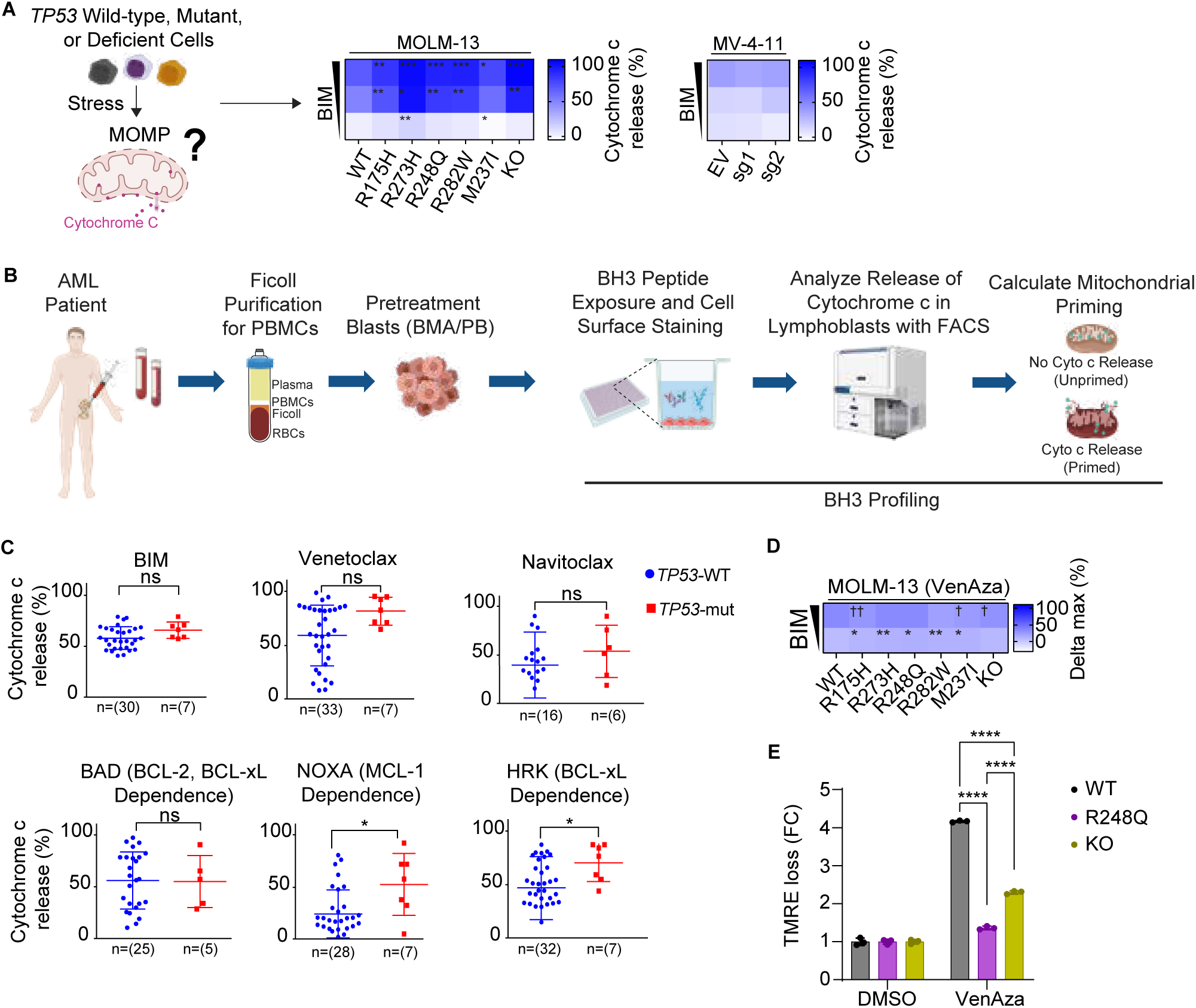
*TP53* mutant AML maintains the potential to undergo MOMP. **(A)** Baseline BH3 profiling heatmap shows mitochondrial priming response of DMSO and titrated BIM for AML (MOLM-13 *TP53*^+/+^*, TP53*^-/mut^, *TP53*^-/-^, and MV-4-11). BIM concentrations used: MOLM-13 *TP53*^+/+^*, TP53*^-/mut^, *TP53*^-/-^: 0.03-3.0 μM; MV-4-11: 0.1-0.5 μM. **(B)** Experimental schematic of the baseline BH3 profiling for primary AML samples. **(C)** BH3 profiling for primary AML samples based on *TP53* status using different peptides and drugs as peptides. **(D)** Dynamic BH3 profiling heatmap after treating MOLM-13 isogenic *TP53^+/+^, TP53^-/mut^*, *TP53^-/-^* cells for 4 hours with VenAza (1 nM venetoclax and 50 nM azacitidine). Dagger = decreased priming. Asterisk = increased priming. **(E)** TMRE loss for *TP53*^+/+^, *TP53*^-/R248Q^, and *TP53^-/-^* cells treated with DMSO or VenAza (venetoclax 3 nM and azacitidine 50 nM) for 24 hours; Analysis via two-way ANOVA followed by Tukey. * p<0.05, ** p<0.01, *** p<0.001, **** p<0.0001.

**Figure 6.**
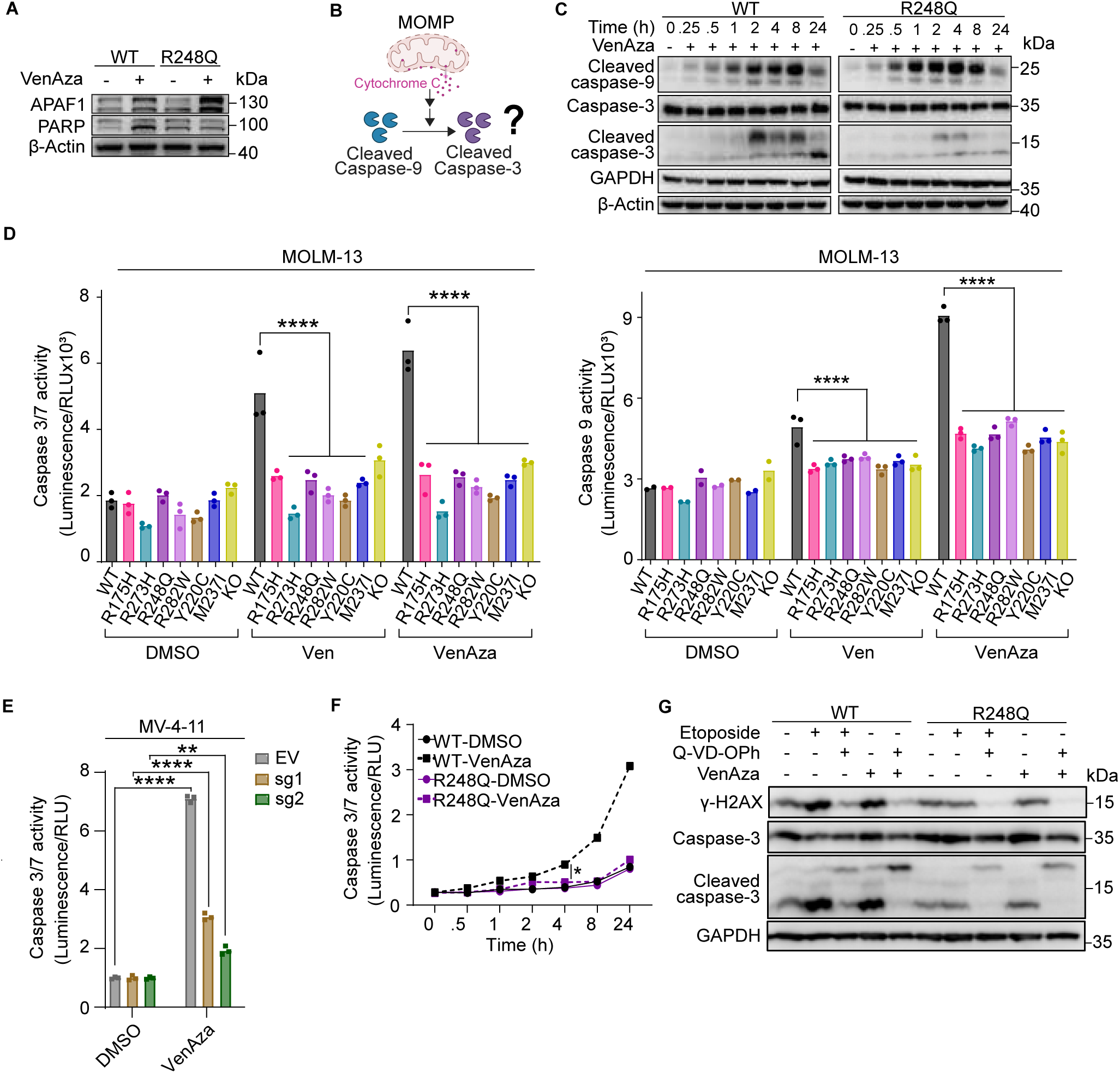
*TP53* mutant AML fails to activate caspases despite MOMP induction in response to therapy. **(A)** Western blot of Apaf-1 and PARP after treatment of MOLM-13 *TP53*^+/+^ and *TP53*^-/R248Q^ with VenAza for 24 hours. **(B)** Schematic question of whether caspase activation occurs after MOMP induction in *TP53* mutant cells. **(C)** Western blot of cleaved caspase-9, cleaved caspase-3 and total caspase-3 after treatment of MOLM-13 *TP53*^+/+^ and *TP53*^-/R248Q^ with VenAza across different timepoints. Both plots developed simultaneously. **(D)** Caspase-3/7 and caspase-9 activity measured in isogenic MOLM-13 *TP53^+/+^, TP53^-/mut^*, *TP53^-/-^* cells after 24 hours of VenAza. Analysis via two-way ANOVA followed by Tukey. **(E)** Caspase-3/7 activity in *TP53*^+/+^ and *TP53*^-/R248Q^ cells treated with DMSO or VenAza (venetoclax 1 nM, azacitidine 50 nM) across 24 hours. **(F)** Western blot of γ-H2AX, caspase-3, and cleaved caspase-3 after pre-treatment with Q-VD-OPh 25 mM for 4 hours and then diluted to 5 mM for 24 hours; treatment with VenAza or etoposide (0.5 uM) for 24 hours. For all figures unless otherwise specified, treatment with VenAza: venetoclax 3 nM, azacitidine 50 nM. * p<0.05, ** p<0.01, *** p<0.001, **** p<0.0001.

To determine whether *TP53* mutant cells resist MOMP following treatment, we performed dynamic BH3 profiling, which showed comparable MOMP induction in *TP53^+/+^*and *TP53^-/mut^* cells after VenAza exposure (Figures 5D). Since mitochondrial membrane potential (MMP) can remain intact when downstream caspase activation is impaired^33,34^, we next measured MMP using tetramethylrhodamine ethyl ester (TMRE) fluorescence. *TP53^-/R248Q^*and *TP53^-/-^* cells had less TMRE fluorescence loss compared to *TP53^+/+^* cells, indicating impaired activation of caspase-9 and caspase-3/7 following VenAza (Figure 5E). Collectively, *TP53* mutant AML cells retain MOMP capacity after direct stress on the mitochondria, and apoptotic failure instead stems from defects of post-mitochondrial permeabilization.

### Impaired caspase activation inhibits drug-induced apoptosis in *TP53* mutant cells

To identify the mechanism that restores apoptotic sensitivity for *TP53* mutant cells, we turned to post-mitochondrial regulators. Apaf-1 was upregulated in response to VenAza in *TP53^+/+^* and *TP53^-/R248Q^* cells while PARP was cleaved only in *TP53^+/+^*(Figure 6A). We then focused on the activation of initiator and executioner caspases to pinpoint where the apoptotic signaling is disrupted (Figure 6B). Time-course analysis following VenAza treatment showed similar cleavage of initiator caspase-9 in *TP53^+/+^* and *TP53^-/R248Q^* cells (Figures 6C and S6A). However, executioner caspase-3 was cleaved selectively in *TP53^+/+^* cells, revealing a disconnect between caspase-9 processing and downstream effector activation in the *TP53* mutant cells. Consistently, caspase-9 and -3/7 enzymatic activity was markedly diminished across all six isogenic *TP53^-/mut^* and *TP53^-/-^* MOLM-13 cells (Figure 6D). Similarly, caspase 3/7 activity was reduced in response to VenAza after deletion of p53 in MV-4-11 cells (Figure 6E). We reasoned that although caspase-9 is cleaved, its enzymatic function is impaired in *TP53* mutant cells, preventing activation of caspase-3 and downstream apoptosis.^35^ Supporting this, VenAza triggered a time-dependent rise in caspase-3/7 activity in *TP53^+/+^*cells but not in *TP53^-/R248Q^* cells (Figure 6F). We also assessed whether reduced caspase-3/7 activation in *TP53^-/R248Q^*and *TP53^-/-^* cells results from defects in the extrinsic apoptotic pathway. Although caspase-8 activity was modestly decreased in these cells, the reduction was far less pronounced than the observed loss for caspase-3/7 activity, suggesting that intrinsic pathway dysfunction is the primary driver of apoptotic failure (Figure S6B).

Given that DNA damage responses play a key role in activating the intrinsic pathway and contribute to drug resistance in *TP53* mutant AML, we next asked whether venetoclax-induced DNA damage is dependent on caspase activation. We pre-treated cells with the pan-caspase inhibitor Q-VD-OPh, followed by VenAza exposure. Etoposide served as a positive control for DNA damage. Caspase inhibition completely abolished γ-H2AX induction in both *TP53^+/+^* and *TP53^-/R248Q^* cells, indicating that DNA damage resulting from VenAza is downstream of caspase activation (Figure 6G). Together, these findings suggest that functional *TP53* is required for full activation of the caspase cascade – particularly the link between caspase-9 and downstream executioner caspases – and is therefore critical for mounting an effective apoptotic response to VenAza in AML.

## DISCUSSION

Therapy resistance remains a formidable barrier to curative treatment in *TP53* mutant cancers, particularly in AML, where patients frequently exhibit primary refractoriness to both conventional chemotherapy and standard of care VenAza.^1–3,8–11^ Clinical and pre-clinical studies have implicated *TP53* loss as a central driver of venetoclax resistance, reinforcing the need for rational combination therapies.^36–38^ Prior studies have shown that *TP53* mutations cause an inferior response to chemotherapy and poor survival in AML and other leukemias. Most have focused on the hypothesis that p53 dysfunction impairs apoptosis through defective transactivation of pro-apoptotic genes. However, our preliminary findings suggest that this model is incomplete.

Canonically, p53 initiates apoptosis by transcriptionally activated BH3-only proteins that trigger mitochondrial outer membrane permeabilization (MOMP), leading to downstream executioner caspase activation and cell death. In *TP53* wild-type AML, transcriptional p53 signaling is robustly activated following VenAza treatment. While this phenomenon is also observed in *TP53* mutant AML, induction of p53 signaling is much lower. This correlates with recent findings in lymphoma, where *TP53* deficient cells lacked feedforward signaling to trigger mitochondrial outer membrane permeabilization (MOMP).^27^ Although the field has acknowledged that MOMP is not always a definitive ‘point-of-no-return,’ its specific relevance to *TP53* mutant tumors has remained unexplored until now. We demonstrate that drug-induced MOMP occurs in leukemia cells harboring *TP53* mutants that abolish transactivation function, suggesting that key regulatory checkpoints exist downstream of MOMP. These findings challenge the prevailing view of p53 as an initiator of apoptosis and instead reveal a previously unrecognized role for functional p53 in sustaining post-MOMP apoptotic signaling.

By redefining the classical ‘point-of-no-return’ in apoptosis, we show that MOMP in not sufficient for cell death in the absence of functional *TP53*. Importantly, this post-MOMP apoptotic blockade was conserved across multiple isogenic *TP53* missense mutants, indicating a shared downstream vulnerability in *TP53* mutant AML. While previous efforts have focused on *TP53* deletions or on reactivating wild-type *TP53* function^17,39^, there is emerging evidence that *TP53* mutations may show distinct phenotypes and clinical outcomes.^40^ Nevertheless, our data suggests that resistance mechanisms persist independently of upstream transcriptional regulation, requiring alternative strategies to restore apoptotic competence.

Collectively, our findings shift the paradigm of apoptotic resistance in *TP53* mutant AML rather than a defect in mitochondrial priming, *TP53* mutant cells have conserved post-mitochondrial failure in caspase activation. This has critical implications for biomarker-driven patient stratification and therapeutic design. Moreover, it raises new mechanistic questions about how *TP53* mutants impair executioner caspase activity – whether via defective apoptosome assembly, altered expression of endogenous caspase inhibitors, or impaired SMAC release. Understanding these mechanisms is essential for developing future therapies that can restore apoptosis and improve outcomes in this high-risk subset.

Ultimately, our work illuminates a previously unrecognized “molecular bottleneck” in apoptosis execution that must be overcome to finally break the therapeutic deadlock in *TP53* mutant AML.

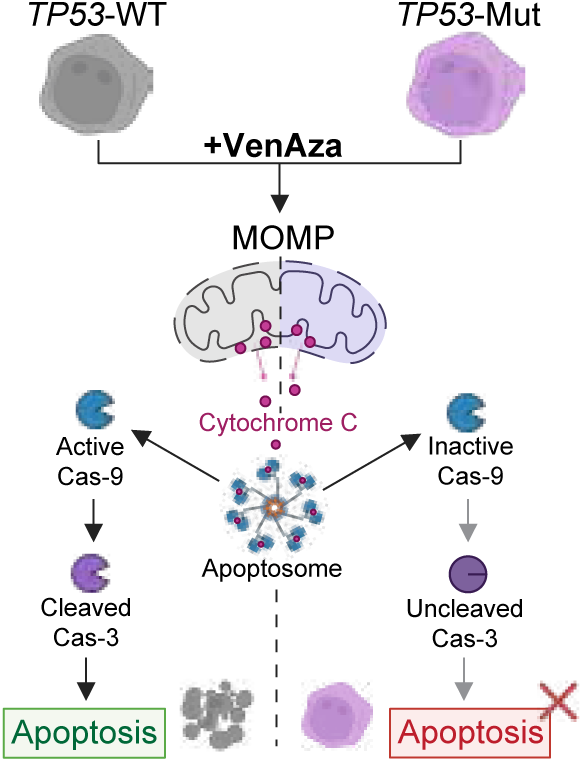

## Acknowledgments

S.B. acknowledges support from the Singapore Ministry of Education (MOE) Academic Research Fund (AcRF) Tier 2 (MOE-000573-00), NUS-Paris University Joint Grant, the Department of Pharmacy and Pharmaceutical Sciences Starup Funds at National University of Singapore. S.B. is a recipient of the EMBO Global Investigator Award. A.M.M. acknowledges support from the Singapore International Graduate Award (SINGA). A.L. acknowledges support from P01 CA066996 and Ludwig Center at Harvard. R.C.L. was supported by a Scholar award from The Leukemia & Lymphoma Society (RCL) and the Edward P. Evans Center for Myelodysplastic Syndrome at Dana-Farber Cancer Institute.

## Contributions

A.M.M. conceptualized and designed experiments, designed figures, performed experiments, analyzed data, and wrote the manuscript. S.J. performed BH3 profiling and DBP experiments and analyzed data. E.A.O. designed figures and wrote the manuscript. F.Q.L. performed experiments, analyzed data, and wrote the manuscript. Y.M. performed validation in isogenic AML cell lines. N.S.E.L., and D.T.E.L. performed experiments and analyzed data. N.C. generated knock-out cells. L.H. designed and performed BH3 profiling experiments on the primary samples. R.C.L. provided samples for BH3 profiling and sequenced *TP53* status from DFCI patients. A.L. supervised and guided BH3 profiling experiments and critically reviewed the manuscript. K.I. critically reviewed the manuscript. S.B. conceptualized the study, designed the experiments, and wrote the manuscript.

## Conflict-of-Interest Disclosure

A.L. owns stocks in Zentalis, Flash Therapeutics, Dialectic, and provided consultancy for Zentalis, Flash Therapeutics, Anji Onco, and Dialectic Therapeutics. R.C.L. provided consultancy for Takeda Pharmaceuticals, Bluebird bio, Qiagen, Sarepta Therapeutics, Verve Therapeutics, Jazz Pharmaceuticals, and Vertex Pharmaceuticals. L.H. owns stocks in and provides consultancy for Pfizer.

---

**Supplemental Figure 1.**
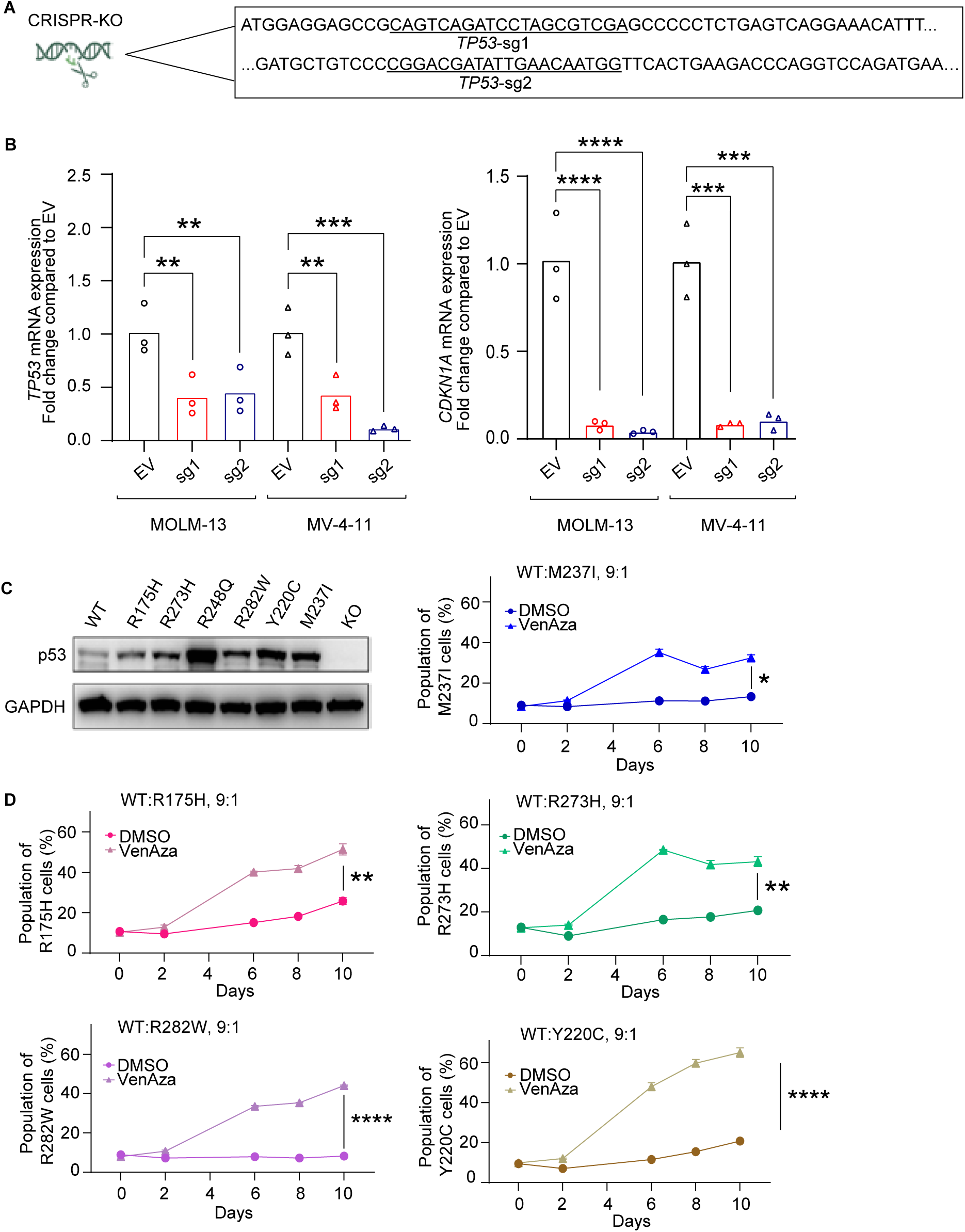
**(A)** Two sgRNAs used for CRISPR knockout. **(B)** Validation of CRISPR *TP53* knockout in isogenic MOLM-13 cells. **(C)** Western blot of p53 expression across *TP53*^+/+^, *TP53*^-/mut^, and *TP53*^-/-^ cells. **(D)** Clonal competition assay evaluating the competition between the cells in coculture after treatment with DMSO or VenAza (venetoclax 3 nM and azacitidine 50 nM) for 10 days.

**Supplemental Figure 2.**
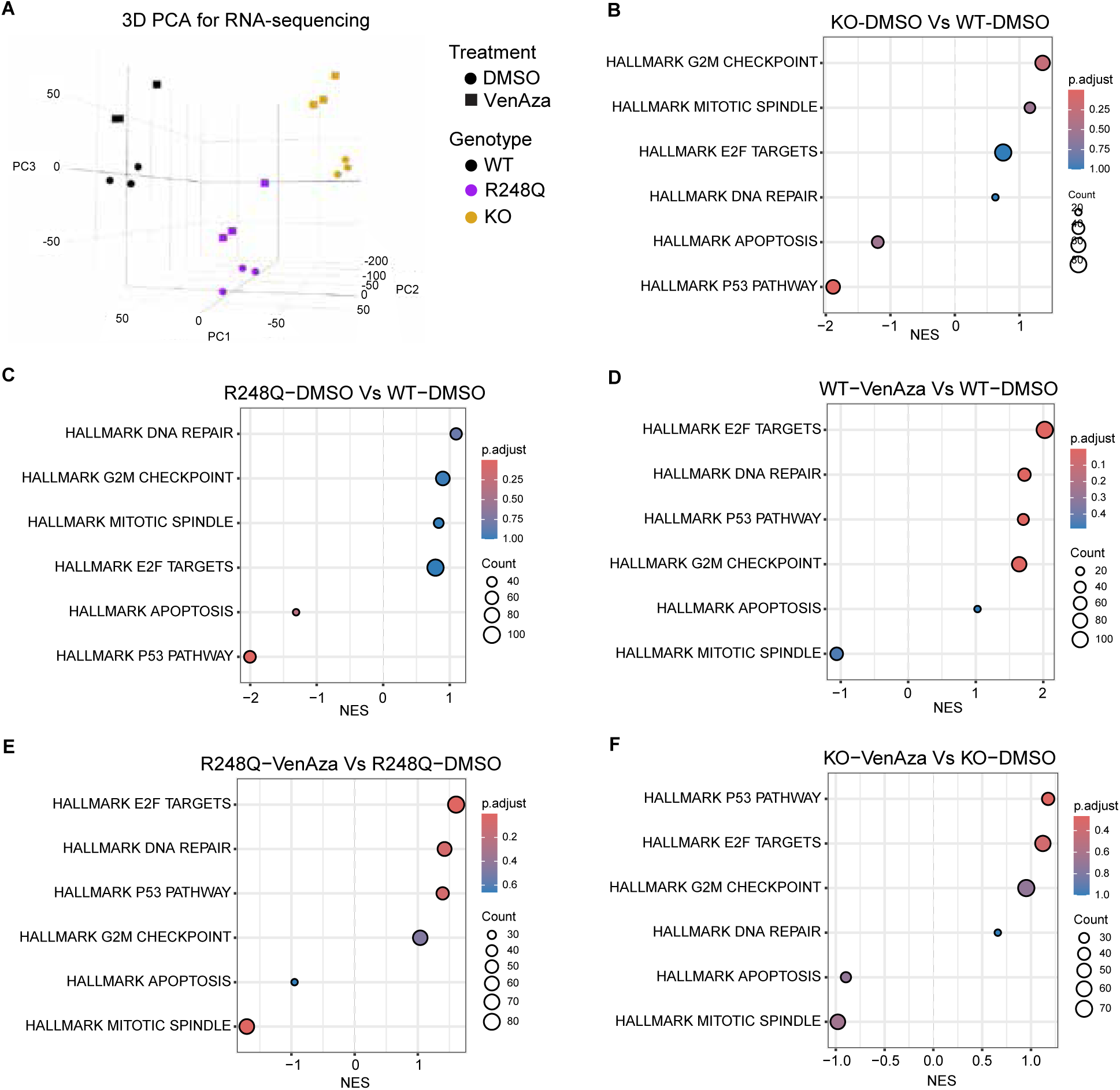
**(A)** Principal component analysis (PCA) of RNA-seq data from all samples. PC1, PC2 and PC3 distinguish samples based on genotype and treatment. Genotypes are color-coded, while treatment conditions are indicated by shape. **(B-F)** Gene set enrichment analysis (GSEA) using MsigDB for pathways: **(B)** *TP53^-/-^* treated with DMSO vs *TP53*^+/+^ treated with DMSO; **(C)** *TP53^-/R248Q^* treated with DMSO vs *TP53*^+/+^ treated with DMSO; **(D)** *TP53^+/+^* treated with VenAza vs *TP53*^+/+^ treated with DMSO; **(E)** *TP53^-/R248Q^* treated with VenAza vs *TP53^-/R248Q^* treated with DMSO; **(F)** *TP53^-/-^* treated with VenAza vs *TP53^-/-^* treated with DMSO. For all figures unless otherwise specified, treatment with VenAza: venetoclax 3 nM, azacitidine 50 nM.

**Supplemental Figure 3.**
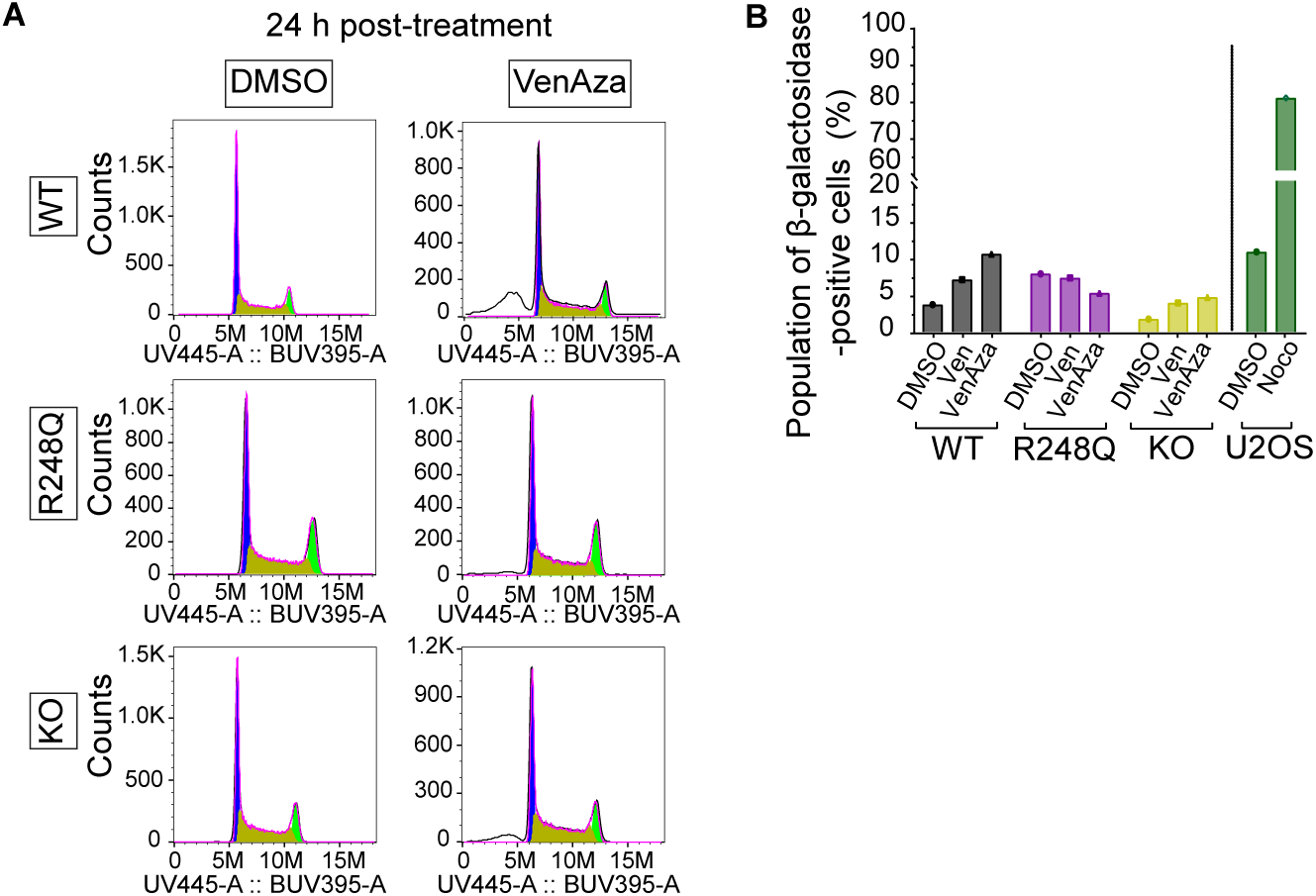
MOLM-13 *TP53*^+/+^, *TP53*^-/R248Q^, and *TP53^-/-^* cells treated with DMSO or VenAza for 24 hours followed by **(A)** cell cycle profiles. **(B)** β-galactosidase assay in *TP53*^+/+^, *TP53*^-/R248Q^, and *TP53*^+/+^ cells treated with DMSO, venetoclax (1 nM), or VenAza (venetoclax 1 nM and azacitidine 50 nM) for 14 days and in U2OS cells treated with DMSO or nocodazole (100 ng/mL) for 4 days.

**Supplemental Figure 4.**
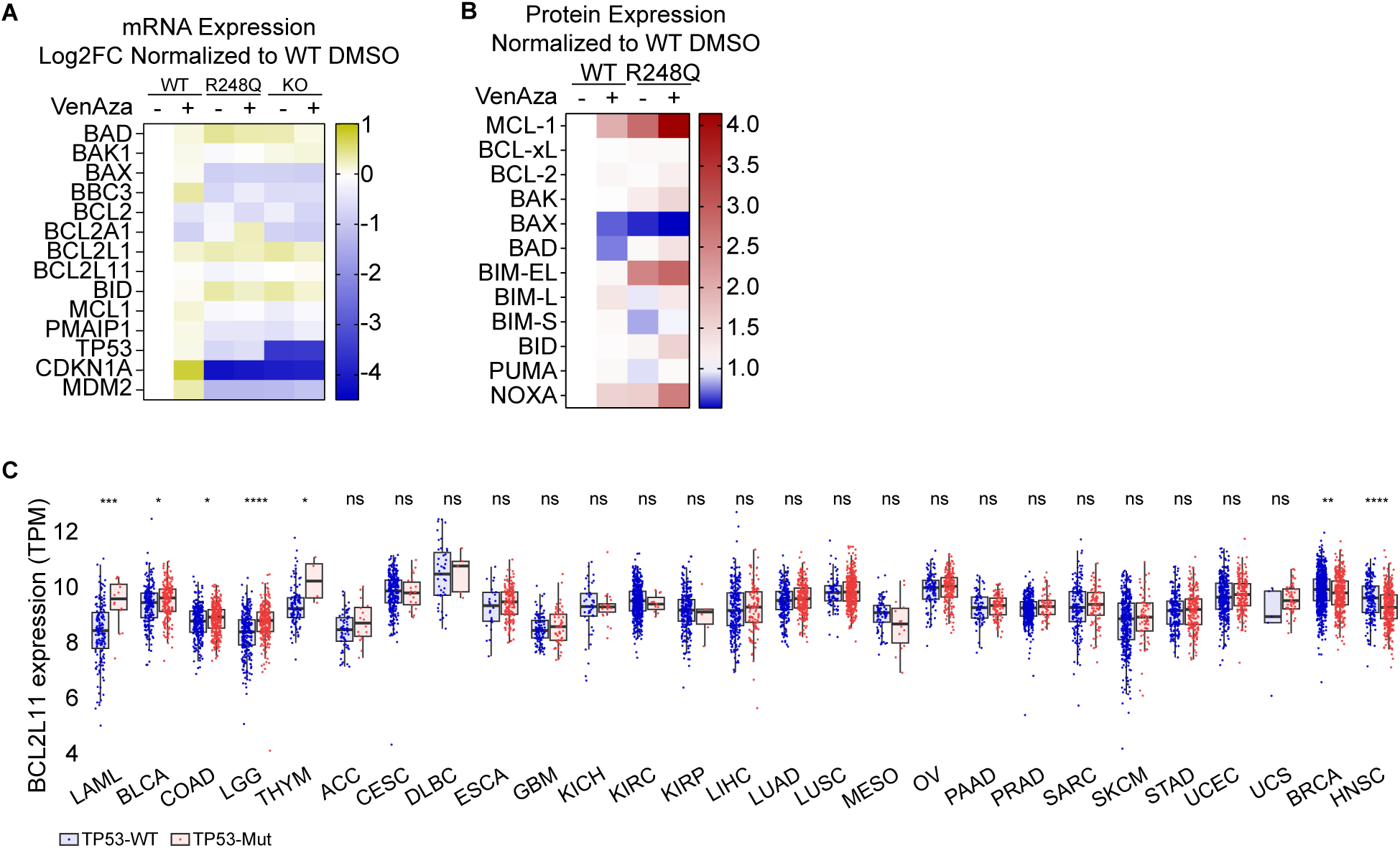
**(A)** Heatmap representing mRNA expression of apoptosis genes obtained from RNA-seq of isogenic MOLM-13 cells normalized to *TP53*^+/+^ cells treated with DMSO. **(B)** Heatmap of densitometry analysis with values normalized to β-Actin or GAPDH, followed by normalization to *TP53*^+/+^ DMSO, corresponding to Figure 4A. **(C)** Heatmap of densitometry analysis with values normalized to β-Actin, followed by normalization to *TP53*^+/+^ DMSO, corresponding to Figure 4B. **(D)** *BCL2L11* mRNA levels across multiple cancers from the TCGA database, stratified according to *TP53* status. Analysis via Wilcoxon test. **(E)** Heatmap of densitometry analysis with values normalized to *TP53*^+/+^, corresponding to Figure 4D.* p<0.05, ** p<0.01, *** p<0.001, **** p<0.0001.

**Supplemental Figure 5.**
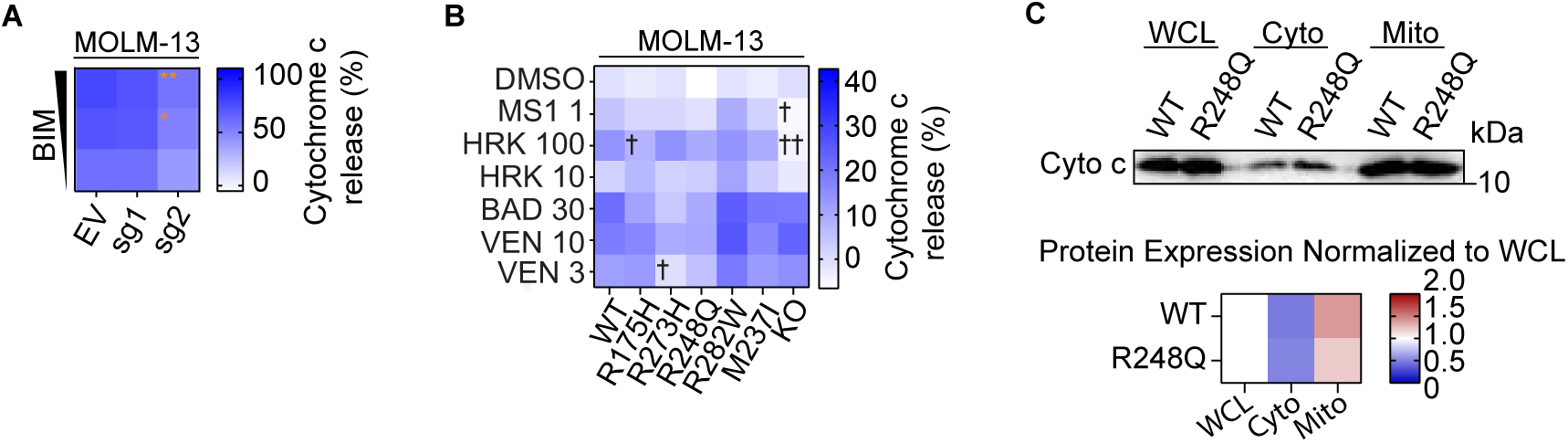
**(A)** Baseline BH3 profiling heatmap shows mitochondrial priming response of DMSO and titrated BIM for MOLM-13 *EV* and *KO* (sg1 and sg2). BIM concentrations used: 0.03-3.0 μM. **(B)** Baseline BH3 profiling heatmap showing the mitochondrial priming response of DMSO, MS-1, HRK, BAD, and venetoclax for MOLM-13 isogenic cells; numbers representing the concentrations in μM. Values normalized to *TP53*^+/+^ DMSO. Dagger = decreased priming. Asterisk = increased priming. **(C)** Western blot of cytochrome c localization in whole cell lysate (WCL), cytoplasm (cyto), and mitochondria (mito). Heatmap of densitometry analysis with values normalized to whole cell lysate (WCL).

**Supplemental Figure 6.**
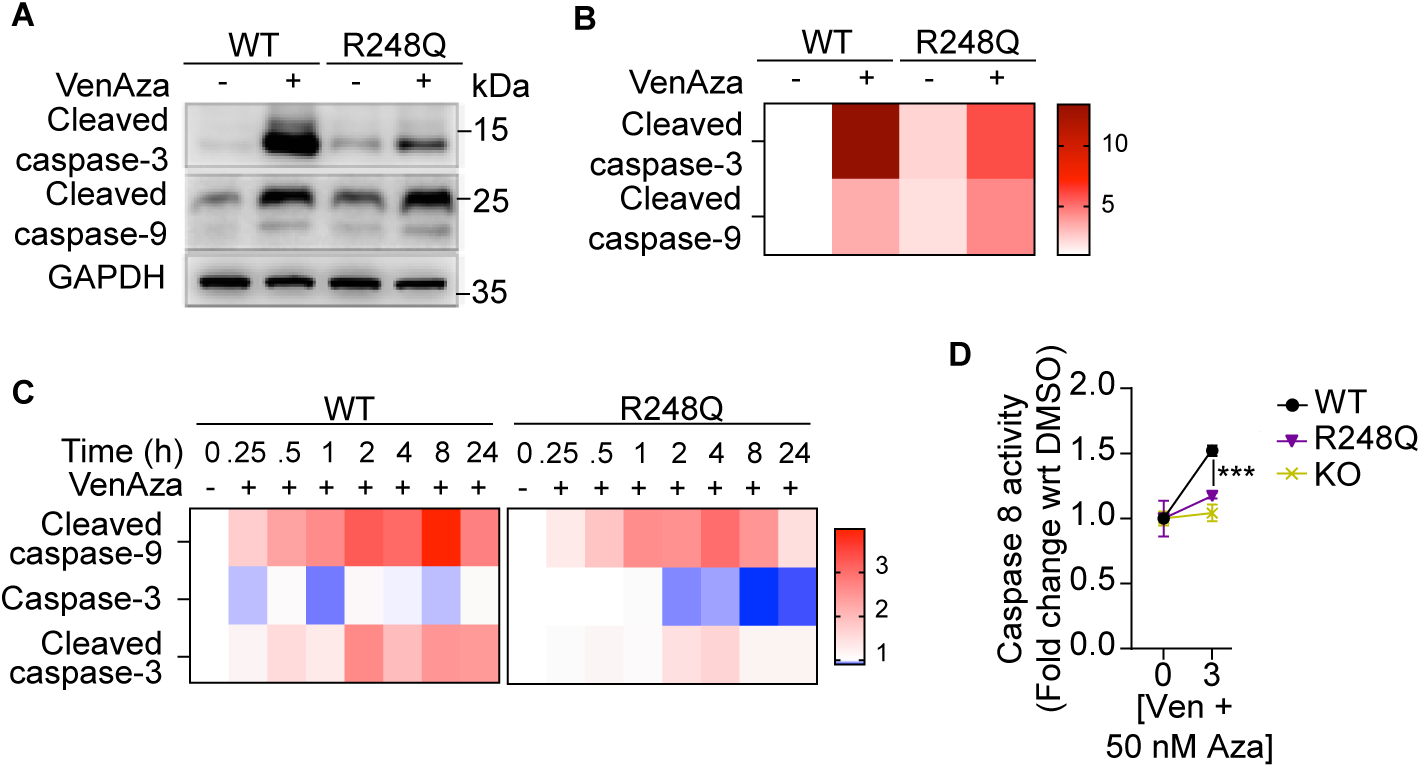
**(A)** Western blot and **(B)** densitometry of cleaved caspase-9, cleaved caspase-3 and total caspase-3 after 24 hours VenAza treatment of MOLM-13 *TP53*^+/+^ and *TP53*^-/R248Q^, run on the same gel. **(C)** Heatmap of densitometry analysis with values normalized to β-Actin or GAPDH, followed by normalization to *TP53*^+/+^ DMSO, corresponding to Figure 6C. **(D)** Caspase-8 activity after treatment with VenAza. For all figures unless otherwise specified, treatment with VenAza: venetoclax 3 nM, azacitidine 50 nM. *** p<0.001.

**Supplemental Table 1:**
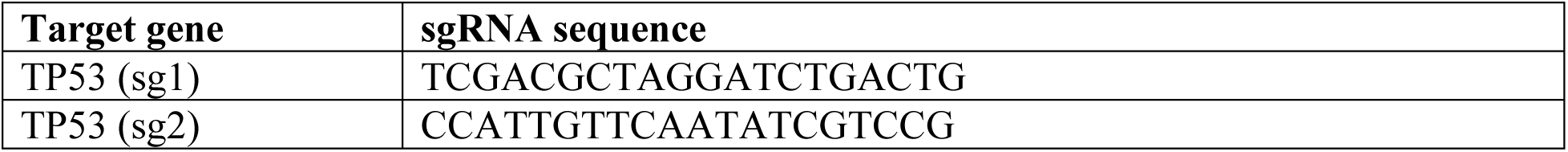
sgRNA sequence used for generation of knock-out cells.

**Supplemental Table 2:** Antibodies used for immunoblotting.

| Antibody | Source | Catalog No. |
| --- | --- | --- |
| p53 Antibody (DO-1) | Santa Cruz Biotechnology | sc-126 |
| Cleaved Caspase-3 (Asp175) Antibody | Cell Signaling Technology | #9661 |
| Cytochrome c Antibody (A-8) | Santa Cruz Biotechnology | sc-13156 |
| Caspase-3 Antibody | Cell Signaling Technology | #9662 |
| GAPDH (D16H11) XP® Rabbit mAb | Cell Signaling Technology | #5174 |
| Bcl-xL (54H6) Rabbit mAb | Cell Signaling Technology | #2764 |
| Mcl-1 (D2W9E) Rabbit mAb | Cell Signaling Technology | #94296 |
| Bcl-2 (124) Mouse mAb | Cell Signaling Technology | #15071 |
| Bak (D4E4) Rabbit mAb | Cell Signaling Technology | #12105 |
| Bax Antibody | Cell Signaling Technology | #2772 |
| Bad (D24A9) Rabbit mAb | Cell Signaling Technology | #9239 |
| Bim (C34C5) Rabbit mAb | Cell Signaling Technology | #2933 |
| BID Antibody | Cell Signaling Technology | #2002 |
| Noxa (D8L7U) Rabbit mAb | Cell Signaling Technology | #14766 |
| Puma (D30C10) Rabbit mAb | Cell Signaling Technology | #12450 |
| Phospho-Histone H2A.X (Ser139) (20E3) Rabbit mAb | Cell Signaling Technology | #9718 |
| Cleaved Caspase-9 (Asp330) (D2D4) Rabbit mAb | Cell Signaling Technology | #7237 |
| β-Actin (8H10D10) Mouse mAb | Cell Signaling Technology | #3700 |
| PARP Antibody | Cell Signaling Technology | #9542 |
| Apaf-1 (R205) Antibody | Cell Signaling Technology | #5088 |

## REFERENCES

1. Toma-Jonik A, Vydra N, Janus P, Widłak W. Interplay between HSF1 and p53 signaling pathways in cancer initiation and progression: non-oncogene and oncogene addiction. Cell Oncol Dordr. 2019;42(5):579–589. doi:10.1007/s13402-019-00452-0

2. Kastenhuber ER, Lowe SW. Putting p53 in Context. Cell. 2017;170(6):1062–1078. doi:10.1016/j.cell.2017.08.028

3. Barbosa K, Li S, Adams PD, Deshpande AJ. The role of TP53 in acute myeloid leukemia: Challenges and opportunities. Genes Chromosomes Cancer. 2019;58(12):875–888. doi:10.1002/gcc.22796

4. Papaemmanuil E, Gerstung M, Bullinger L, et al. Genomic Classification and Prognosis in Acute Myeloid Leukemia. N Engl J Med. 2016;374(23):2209–2221. doi:10.1056/NEJMoa1516192

5. Lindsley RC, Saber W, Mar BG, et al. Prognostic Mutations in Myelodysplastic Syndrome after Stem-Cell Transplantation. N Engl J Med. 2017;376(6):536–547. doi:10.1056/NEJMoa1611604

6. Feurstein S, Rücker FG, Bullinger L, et al. Haploinsufficiency of ETV6 and CDKN1B in patients with acute myeloid leukemia and complex karyotype. BMC Genomics. 2014;15(1):784. doi:10.1186/1471-2164-15-784

7. Pollyea DA, Pratz KW, Wei AH, et al. Outcomes in Patients with Poor-Risk Cytogenetics with or without TP53 Mutations Treated with Venetoclax and Azacitidine. Clin Cancer Res Off J Am Assoc Cancer Res. 2022;28(24):5272–5279. doi:10.1158/1078-0432.CCR-22-1183

8. DiNardo CD, Pratz K, Pullarkat V, et al. Venetoclax combined with decitabine or azacitidine in treatment-naive, elderly patients with acute myeloid leukemia. Blood. 2019;133(1):7–17. doi:10.1182/blood-2018-08-868752

9. Konopleva M, Letai A. BCL-2 inhibition in AML: an unexpected bonus? Blood. 2018;132(10):1007–1012. doi:10.1182/blood-2018-03-828269

10. Wei AH, Strickland SA, Hou JZ, et al. Venetoclax Combined With Low-Dose Cytarabine for Previously Untreated Patients With Acute Myeloid Leukemia: Results From a Phase Ib/II Study. J Clin Oncol Off J Am Soc Clin Oncol. 2019;37(15):1277–1284. doi:10.1200/JCO.18.01600

11. DiNardo CD, Jonas BA, Pullarkat V, et al. Azacitidine and Venetoclax in Previously Untreated Acute Myeloid Leukemia. N Engl J Med. 2020;383(7):617–629. doi:10.1056/NEJMoa2012971

12. Kim K, Maiti A, Loghavi S, et al. Outcomes of TP53-mutant acute myeloid leukemia with decitabine and venetoclax. Cancer. 2021;127(20):3772–3781. doi:10.1002/cncr.33689

13. Daver N, Wei AH, Pollyea DA, Fathi AT, Vyas P, DiNardo CD. New directions for emerging therapies in acute myeloid leukemia: the next chapter. Blood Cancer J. 2020;10(10):107. doi:10.1038/s41408-020-00376-1

14. Green DR, Kroemer G. Cytoplasmic functions of the tumour suppressor p53. Nature. 2009;458(7242):1127-1130. doi:10.1038/nature07986

15. Yu J, Wang Z, Kinzler KW, Vogelstein B, Zhang L. PUMA mediates the apoptotic response to p53 in colorectal cancer cells. Proc Natl Acad Sci U S A. 2003;100(4):1931–1936. doi:10.1073/pnas.2627984100

16. Oda E, Ohki R, Murasawa H, et al. Noxa, a BH3-only member of the Bcl-2 family and candidate mediator of p53-induced apoptosis. Science. 2000;288(5468):1053–1058. doi:10.1126/science.288.5468.1053

17. Thijssen R, Diepstraten ST, Moujalled D, et al. Intact TP-53 function is essential for sustaining durable responses to BH3-mimetic drugs in leukemias. Blood. 2021;137(20):2721–2735. doi:10.1182/blood.2020010167

18. Miyashita T, Harigai M, Hanada M, Reed JC. Identification of a p53-dependent negative response element in the bcl-2 gene. Cancer Res. 1994;54(12):3131–3135.

19. Wu Y, Mehew JW, Heckman CA, Arcinas M, Boxer LM. Negative regulation of bcl-2 expression by p53 in hematopoietic cells. Oncogene. 2001;20(2):240–251. doi:10.1038/sj.onc.1204067

20. Chipuk JE, Kuwana T, Bouchier-Hayes L, et al. Direct activation of Bax by p53 mediates mitochondrial membrane permeabilization and apoptosis. Science. 2004;303(5660):1010–1014. doi:10.1126/science.1092734

21. Chipuk JE, Bouchier-Hayes L, Kuwana T, Newmeyer DD, Green DR. PUMA couples the nuclear and cytoplasmic proapoptotic function of p53. Science. 2005;309(5741):1732–1735. doi:10.1126/science.1114297

22. Leu JIJ, Dumont P, Hafey M, Murphy ME, George DL. Mitochondrial p53 activates Bak and causes disruption of a Bak-Mcl1 complex. Nat Cell Biol. 2004;6(5):443–450. doi:10.1038/ncb1123

23. Tait SWG, Parsons MJ, Llambi F, et al. Resistance to caspase-independent cell death requires persistence of intact mitochondria. Dev Cell. 2010;18(5):802–813. doi:10.1016/j.devcel.2010.03.014

24. Ryan J, Montero J, Rocco J, Letai A. iBH3: simple, fixable BH3 profiling to determine apoptotic priming in primary tissue by flow cytometry. Biol Chem. 2016;397(7):671–678. doi:10.1515/hsz-2016-0107

25. Boettcher S, Miller PG, Sharma R, et al. A dominant-negative effect drives selection of TP53 missense mutations in myeloid malignancies. Science. 2019;365(6453):599–604. doi:10.1126/science.aax3649

26. Schimmer RR, Kovtonyuk LV, Klemm N, et al. TP53 mutations confer resistance to hypomethylating agents and BCL-2 inhibition in myeloid neoplasms. Blood Adv. 2022;6(11):3201–3206. doi:10.1182/bloodadvances.2021005859

27. Diepstraten ST, Yuan Y, La Marca JE, et al. Putting the STING back into BH3-mimetic drugs for TP53-mutant blood cancers. Cancer Cell. 2024;42(5):850–868.e9. doi:10.1016/j.ccell.2024.04.004

28. Bhatt S, Pioso MS, Olesinski EA, et al. Reduced Mitochondrial Apoptotic Priming Drives Resistance to BH3 Mimetics in Acute Myeloid Leukemia. Cancer Cell. 2020;38(6):872–890.e6. doi:10.1016/j.ccell.2020.10.010

29. Zhang Q, Riley-Gillis B, Han L, et al. Activation of RAS/MAPK pathway confers MCL-1 mediated acquired resistance to BCL-2 inhibitor venetoclax in acute myeloid leukemia. Signal Transduct Target Ther. 2022;7(1):1–13. doi:10.1038/s41392-021-00870-3

30. Niu X, Zhao J, Ma J, et al. Binding of Released Bim to Mcl-1 is a Mechanism of Intrinsic Resistance to ABT-199 which can be Overcome by Combination with Daunorubicin or Cytarabine in AML Cells. Clin Cancer Res Off J Am Assoc Cancer Res. 2016;22(17):4440–4451. doi:10.1158/1078-0432.CCR-15-3057

31. Carter BZ, Mak PY, Tao W, et al. Combined inhibition of BCL-2 and MCL-1 overcomes BAX deficiency-mediated resistance of TP53-mutant acute myeloid leukemia to individual BH3 mimetics. Blood Cancer J. 2023;13(1):1–13. doi:10.1038/s41408-023-00830-w

32. Yuda J, Will C, Phillips DC, et al. Selective MCL-1 inhibitor ABBV-467 is efficacious in tumor models but is associated with cardiac troponin increases in patients. Commun Med. 2023;3(1):154. doi:10.1038/s43856-023-00380-z

33. Cepero E, King AM, Coffey LM, Perez RG, Boise LH. Caspase-9 and effector caspases have sequential and distinct effects on mitochondria. Oncogene. 2005;24(42):6354–6366. doi:10.1038/sj.onc.1208793

34. Ricci JE, Gottlieb RA, Green DR. Caspase-mediated loss of mitochondrial function and generation of reactive oxygen species during apoptosis. J Cell Biol. 2003;160(1):65–75. doi:10.1083/jcb.200208089

35. Bratton SB, Salvesen GS. Regulation of the Apaf-1–caspase-9 apoptosome. J Cell Sci. 2010;123(19):3209. doi:10.1242/jcs.073643

36. Chen X, Glytsou C, Zhou H, et al. Targeting Mitochondrial Structure Sensitizes Acute Myeloid Leukemia to Venetoclax Treatment. Cancer Discov. 2019;9(7):890–909. doi:10.1158/2159-8290.CD-19-0117

37. Nechiporuk T, Kurtz SE, Nikolova O, et al. The TP53 Apoptotic Network Is a Primary Mediator of Resistance to BCL2 Inhibition in AML Cells. Cancer Discov. 2019;9(7):910–925. doi:10.1158/2159-8290.CD-19-0125

38. Savona MR, Rathmell JC. Mitochondrial Homeostasis in AML and Gasping for Response in Resistance to BCL2 Blockade. Cancer Discov. 2019;9(7):831–833. doi:10.1158/2159-8290.CD-19-0510

39. Nishida Y, Ishizawa J, Ayoub E, et al. Enhanced TP53 reactivation disrupts MYC transcriptional program and overcomes venetoclax resistance in acute myeloid leukemias. Sci Adv. 2023;9(48):eadh1436. doi:10.1126/sciadv.adh1436

40. Hassin O, Nataraj NB, Shreberk-Shaked M, et al. Different hotspot p53 mutants exert distinct phenotypes and predict outcome of colorectal cancer patients. Nat Commun. 2022;13(1):2800. doi:10.1038/s41467-022-30481-7

